# Multi-modal analysis and integration of single-cell morphological data

**DOI:** 10.1101/2022.05.19.492525

**Authors:** Kiya W. Govek, Jake Crawford, Artur B. Saturnino, Kristi Zoga, Michael P. Hart, Pablo G. Camara

## Abstract

High-resolution imaging-based single-cell profiling has transformed the study of cells in their spatial context. However, the lack of quantitative methods that can summarize the great diversity of complex cell shapes found in tissues and infer associations with other single-cell data modalities limits current analyses. Here, we report a general computational framework for the multi-modal analysis and integration of single-cell morphological data. We build upon metric geometry to construct cell morphology latent spaces, where distances in these spaces indicate the amount of physical deformation needed to change the morphology of one cell into that of another. Using these spaces, we integrate morphological data across technologies and leverage associated single-cell RNA-seq data to infer relations between morphological and transcriptomic cellular processes. We apply this framework to imaging and multi-modal data of neurons and glia to uncover genes related to neuronal plasticity. Our approach represents a strategy for incorporating cell morphological data into single-cell omics analyses.

## Introduction

Since the advent of staining techniques in the 19^th^ century, cell morphology has become one of the most described phenotypes in biology. The idea that the morphology of a cell is related to its function has been central to major discoveries, such as the neuron doctrine (Ramón y Cajal, 1960), the molecular basis of sickle cell disease (Pauling et al., 1949), and the pathways for cell migration and chemotropic sensing (Wessells et al., 1971). In the nervous system, whole-cell patch-clamp has enabled the morphological reconstruction and electrophysiological recording of >100,000 neurons (Ascoli et al., 2007). The incorporation of high-throughput single-cell RNA-seq onto this technique, known as Patch-seq (Bardy et al., 2016; Cadwell et al., 2016; Chen et al., 2016; Foldy et al., 2016; Fuzik et al., 2016), has opened the door to deeper characterizations that include morphological, transcriptomic, and electrophysiological information from the same cells (Lipovsek et al., 2021). More broadly, the recent explosion of spatially resolved technologies for single-cell transcriptomics has transformed the study of cells in their spatial context by enabling researchers to infer links between morphological and molecular phenotypes (Asp et al., 2020; Liao et al., 2021; Moses and Pachter, 2022). The potential of this new array of techniques is immense, not only for cell taxonomic purposes, but also for uncovering the pathways that are associated with, and may ultimately drive, the morphological diversification and plasticity of cells.

Algorithms for cell morphometry seek to determine similarities among the morphology of individual cells in digitally reconstructed microscopy images. Current methods extract a set of shape descriptors that summarize the morphology of each cell. Simple geometric descriptors like the area and perimeter of the cell (Carpenter et al., 2006; Legland et al., 2016) can be applied to most cell types, but have limited power to accurately discriminate complex cell morphologies like those of neurons and glia. On the other hand, more complex cell-type-specific descriptors, such as neuronal branching patterns (Arshadi et al., 2021; Scorcioni et al., 2008; Sholl, 1953), need to be tailored to the specific cell type of interest and cannot be applied broadly. In addition, they are arbitrary with respect to the features that are used, and the weight assigned to them. To overcome these limitations, other methods introduce similarity scores based on tree alignment (Costa et al., 2016; Wan et al., 2015) or decomposition in Fourier, Zernike, or spherical harmonic moments (Brechbühler et al., 1995; Khotanzad and Hong, 1990; Medyukhina et al., 2020; Pincus and Theriot, 2007). However, these methods require building combinations of descriptors that are invariant under rigid transformations or carefully pre-aligning the cells using Procrustes analysis, and fail to quantify morphological differences between highly dissimilar cells. In general, none of the current approaches reflect the biophysical processes involved in cell morphological changes or lead to an actual distance function in the cell morphology space. These limitations have precluded the development of advanced algebraic and statistical approaches for the analysis of cell morphological data, such as methods for integrating these data, constructing consensus cell morphologies, or inferring cell state trajectories associated with morphological processes.

Here, we build upon recent developments in applied metric geometry and shape registration (Mémoli, 2007, 2011; Mémoli and Sapiro, 2005) to establish a general computational framework for studying complex and heterogeneous cell morphologies across the broad range of cells found in tissues. This framework enables the characterization of morphological cellular processes from a biophysical perspective and produces an actual mathematical distance upon which rigorous algebraic and statistical analytic approaches can be built. The approach has the generality and stability of simple geometric shape descriptors, the discriminative power of cell-type specific descriptors, and the unbiasedness and hierarchical structure of moments-based descriptors. Using this framework, we address several outstanding methodological roadblocks in relating cell morphology to molecular content and function, including the combined analysis of morphological, molecular, and physiological information of individual cells, and the integration of morphological information across experiments and technologies. Applying this framework to Patch-seq, patch-clamp, electron, and two-photon microscopy data of neurons and glia, we identify some of the genetic and molecular programs that are associated with the plasticity of neurons. Taken together, the results of these analyses demonstrate that the application of metric geometry to the study of cell morphology not only greatly increases the accuracy and versatility of cell morphology analyses, but also enables currently unavailable analyses such as the integration of cell morphological data with other single-cell data modalities and across experiments. We have implemented this analytic framework in an open-source software, which we expect will be useful to researchers working with single-cell morphological data of any kind.

## Results

### A general framework for the quantitative analysis of cell morphology data

In its simplest formulation, the study of cell morphology involves the quantitative comparison of cell shapes irrespective of distance-preserving transformations (*isometries*), such as rotations and translations. From a mathematical standpoint, this is a problem of metric geometry. The Gromov-Hausdorf (GH) distance (Edwards, 1975; Gromov, 1981) measures how far two compact metric spaces are from being isometric. In physical terms, it determines the minimum amount of deformation required to convert the shape of an object into that of another. The use of the GH distance to describe cell shapes is therefore broadly applicable to any cell type, as it does not rely on geometric features that are particular to the cell type or require pre-aligning the cells to a reference shape. Because of these reasons, we sought to develop a general framework for cell morphometry by building upon these concepts in metric geometry. Since computing the GH distance is intractable even for relatively small datasets, we based our approach upon a computationally efficient approximation, referred to as the Gromov-Wasserstein (GW) distance (Mémoli, 2007, 2011; Mémoli and Sapiro, 2005). The GW distance preserves most of the mathematical properties of the GH distance and leads to an actual distance function (Mémoli, 2011). Although its running time grows cubically with the number of points, its efficiency can be further improved by means of nearly linear-time approximations that build upon optimal transport regularization (Scetbon et al., 2021; Solomon et al., 2016) and nesting strategies (Chowdhury et al., 2021).

The starting point to our analytic framework is the 2D segmentation masks or 3D digital reconstructions of individual cells, which are discretized by evenly sampling points from their outline (Fig. 1A). For each cell, we compute the pairwise distance matrix (*d_i_*) between its sampled points. Then, for each pair of cells, *i* and *j*, we compute the GW distance between the matrices *d_i_* and *d_j_* using optimal transport (Fig. 1B). The result is a pairwise GW distance that quantifies the morphological differences between each pair of cells.

**Figure 1.**
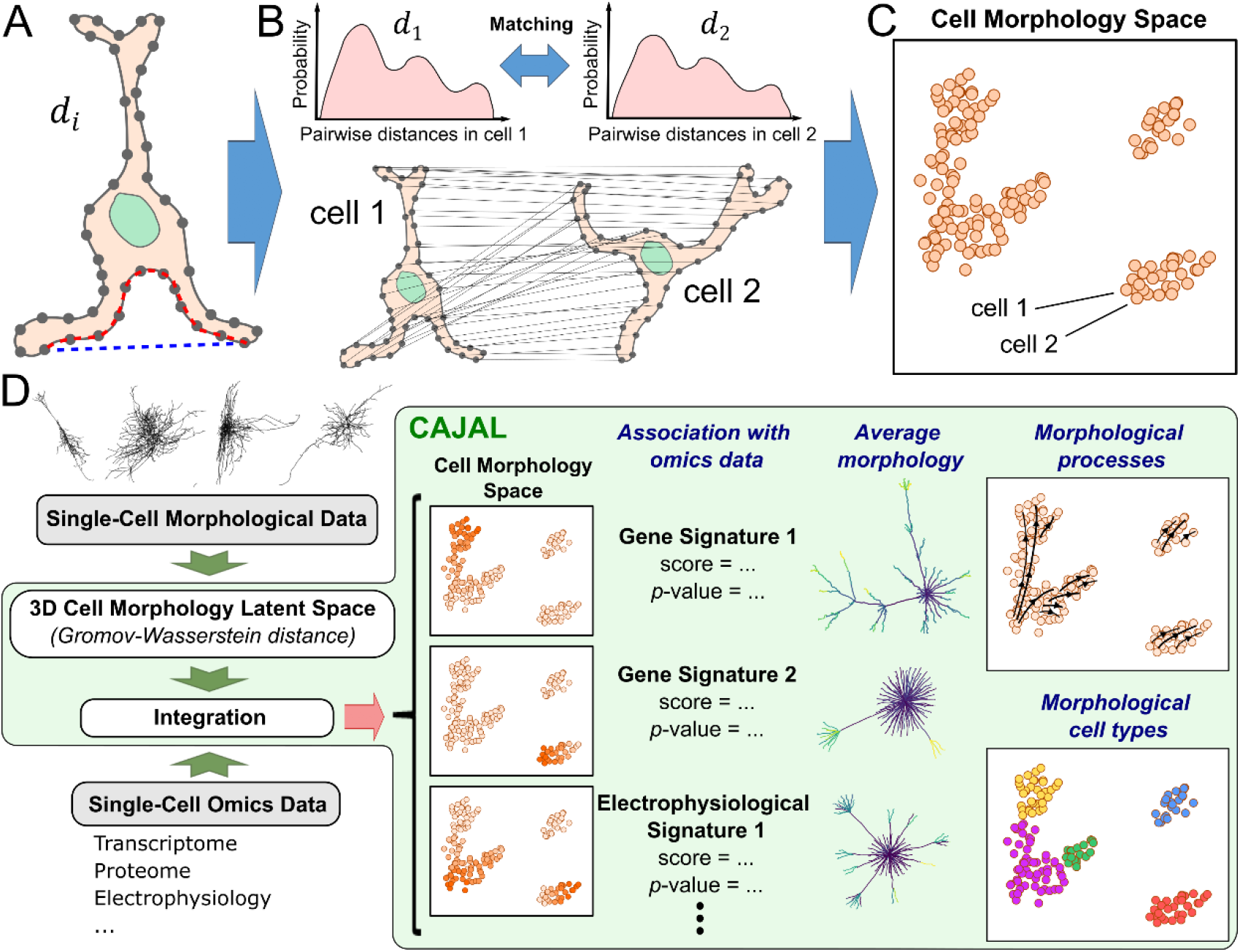
A general framework for the quantitative analysis and integration of cell morphological data based on metric geometry. **A)** *CAJAL* takes as input the 2D or 3D cell segmentation masks or traces of a set of cells. For each cell, a set of points is evenly sampled from the outline and their pairwise distance matrix *d_i_* is computed. The Euclidean and geodesic distances between 2 sampled points is indicated with a blue and red dashed line, respectively. Different metrics for computing *d_i_* capture different aspects of cell morphology. **B)** An optimal matching between the discretized morphologies of each pair of cells is established by computing the GW distance between their corresponding *d_i_* matrices. Computationally efficient approximations to the GW distance use optimal transport theory to establish a map between the distributions of sampled points pairwise distances in each cell. The value of the cost function at the minimum measures the amount of deformation that is needed to convert the shape of one cell in that of another. **C)** The GW distance matrix can be thought of as a distance in a latent space of cell morphologies. This enables the application of statistical and machine learning methods for the analysis and integration of point clouds. **D)** Overview of the open-source software *CAJAL*. The software takes single-cell morphological data in the form of segmentation masks or neuronal traces as input and enables its integration with other single-cell data modalities, the visualization and clustering of cell morphology spaces, the identification of molecular and electrophysiological features associated with changes in cell morphology, the computation of average or representative cell shapes, and the visualization of trajectories in the cell morphology space.

Different metrics for measuring distances between sampled points lead to different properties of the GW distance that may be advantageous in specific applications (Fig. 1A). For example, using Euclidean distance results in a GW matrix that accounts for the positioning of cell appendages, which can be particularly useful in the study of neuronal projections. On the other hand, using geodesic distance results in a GW matrix that is invariant under bending deformations of the cell, and it is therefore particularly sensitive to topological features such as the branching structure of cell appendages.

In all cases, the resulting GW distance can be thought of as a distance in a latent space of cell morphologies (Fig. 1C). In this latent space, each cell is represented by a point, and distances between cells indicate the amount of physical deformation needed to change the morphology of one cell into that of another. By formulating the problem in this way, we can use statistical and machine learning approaches to define cell populations based on their morphology; dimensionally reduce and visualize cell morphology spaces; integrate cell morphology spaces across tissues, technologies, and with other single-cell data modalities (for example, single-cell RNA-seq or ATAC-seq data); or infer trajectories associated with continuous morphological processes. We have implemented these analyses in an open-source Python library, called *CAJAL*, which can be used with arbitrarily complex and heterogeneous cell populations (Fig. 1D).

### GW cell morphology spaces accurately summarize complex cell shapes

To assess the ability of GW cell morphology spaces to summarize complex cell shapes, we applied *CAJAL* to the 3D basal and apical dendrite reconstructions of 506 neurons from the mouse visual cortex profiled with patch-clamp (Gouwens et al., 2019). The resulting space of cell morphologies recapitulated the neuronal morphological types of the visual cortex (Fig. 2A). Cells with a similar morphology appeared in proximity in the UMAP representation of this space. Molecularly defined neuronal types were also localized in the representation (Fig. 2B), consistent with the presence of morphological characteristics that are unique to each molecular subtype. Excitatory and inhibitory neurons clustered separately, and individual neurons were organized in the cell morphology space according to their cortical layer and Cre driver line (Fig. 2B). Clustering the morphology space using Louvain community detection (Blondel et al., 2008) partitioned it into 9 morphological populations. Using the metric structure of the cell morphology space, we then computed the medoid and average cell morphology for each cluster (Fig. 2C). These summaries accurately represented the main morphological characteristics of the cell populations and were consistent with the diversity of neuronal morphologies found in the mouse visual cortex (Gouwens et al., 2019).

**Figure 2.**
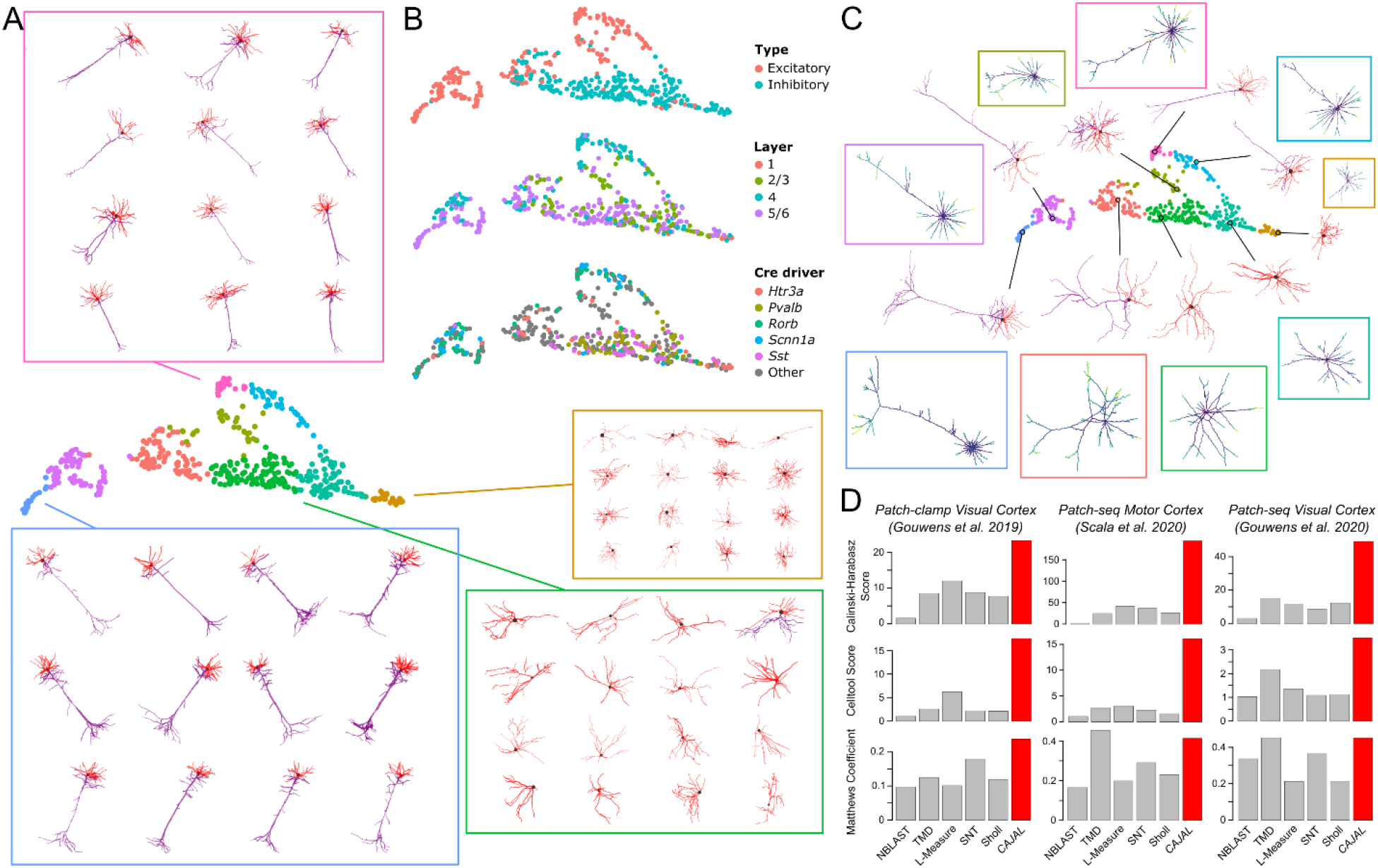
Cell morphology spaces accurately summarize the complexity of cell shapes. **A)** UMAP representation of the cell morphology space of 506 neurons from the murine visual cortex profiled with whole-cell patch-clamp. The representation is colored by the morphological cell populations that resulted from clustering the cell morphology space using Louvain community detection. The morphologies of individual neurons sampled from 4 of the populations are shown for reference. Apical and basal dendrites are indicated in purple and red, respectively. **B)** The UMAP representation is colored by the neuronal type (top), cortical layer (middle), and Cre driver line (bottom). The GW cell morphology space captures morphological differences between neurons of different molecular type and anatomic location. **C)** The metric structure of the GW morphology space enables performing algebraic operations such as averaging shapes. The figure shows the medoid (indicated with a circle) and average morphologies (in boxes) computed for each of the morphological populations in (A). **D)** The ability of *CAJAL* to identify morphological differences between molecularly defined neurons is assessed in 3 datasets in comparison to 5 currently available methods. In this study, *CAJAL* offered substantially better or similar results compared to the best performing method in each dataset according to the three metrics that were used for the evaluation.

To quantitatively evaluate the ability of the GW distance to accurately summarize complex neuron morphologies in comparison to state-of-the-art methods for neuron morphometry, we considered two Patch-seq datasets of the visual (Gouwens et al., 2020) and motor cortex (Scala et al., 2021) in addition to the patch-clamp dataset of the visual cortex. For each dataset, we assessed the ability of *CAJAL* and 5 other methods (Sholl analysis (Sholl, 1953), L-measure (Scorcioni et al., 2008), SNT (Arshadi et al., 2021), NBLAST (Costa et al., 2016), and TMD (Kanari et al., 2018)) to identify morphological differences between molecularly-defined neurons. In the case of the patch-clamp dataset, we considered neurons labeled with different Cre driver lines, for a total of 31 lines, with the understanding that each line preferentially labels distinct neuronal types. In the case of the two Patch-seq datasets, we considered the known classification of motor and visual cortex neurons into 9 and 6 transcriptionally-defined subtypes (t-types), respectively, based on their gene expression profile (Gouwens et al., 2020; Scala et al., 2021). We used three different metrics of performance to evaluate the ability of each method to predict the molecular type of each individual neuron based on its morphology: the multi-class Matthews Correlation Coefficient of a *k* nearest neighbor classifier, the Calinski-Harabasz clustering score of neurons from the same molecular type in the cell morphology space, and the benchmarking score introduced in a previous study of cell shape analysis methods (Pincus and Theriot, 2007). In this comparative study, we found that the GW distance outperformed existing methods for neuron morphometry (Fig. 2D). In addition, some of the best performing methods, such as TMD, produced errors and were not able to summarize the morphology of 26 neurons for which the model assumptions were violated. A similar evaluation of the ability of each algorithm to identify 47 t-types of inhibitory neurons and 26 t-types of excitatory neurons in the motor cortex (Scala et al., 2021) led to consistent results (Supplementary Fig. 1). As expected, the accuracy and running time of *CAJAL* in these analyses increased with the number of points that are sampled from the outline of the cell (Supplementary Fig. 2). However, for these datasets the accuracy saturated at approximately 100 points, indicating no major advantage in sampling a larger number of points. Taken together, these results demonstrate the utility and versatility of the GW distance to perform unbiased studies of complex cell morphologies.

### GW cell morphology spaces recapitulate heterogeneous cell types

We next evaluated the ability of the GW distance to summarize cell morphologies across heterogenous cell populations. For that purpose, we used *CAJAL* to study the morphologies of 70,510 cells from a cubic millimeter volume of the mouse visual cortex profiled by the Machine Intelligence from Cortical Networks (MICrONS) program using two-photon microscopy, microtomography, and serial electron microscopy (MICrONS Consortium et al., 2021). This dataset includes not only neurons, but also multiple types of glia and immune cells. The UMAP representation of the cell morphology space produced by *CAJAL* recapitulated in an unsupervised manner the broad spectrum of cell types that are present in the tissue, including several populations of neurons, astrocytes, microglia, and immune cells (Fig. 3A). These populations were consistent with the manual annotations of 185 individual cells provided by the MICrONS program (Fig. 3B). Neurons from different cortical layers were associated with distinct regions of the cell morphology space, indicating the presence of morphological differences between neurons from different layers (Fig. 3C). In addition, our analysis uncovered morphological differences between astrocytes located in different cortical layers (Figs. 3D, E). Specifically, layer 1 astrocytes were substantially smaller, and layer 2/3 astrocytes were elongated perpendicularly to the pial surface, in comparison to astrocytes residing in other cortical layers (Figs. 3D, E). These morphological differences were consistent with recent observations in Glast-EMTB-GFP transgenic mice (Lanjakornsiripan et al., 2018).

**Figure 3.**
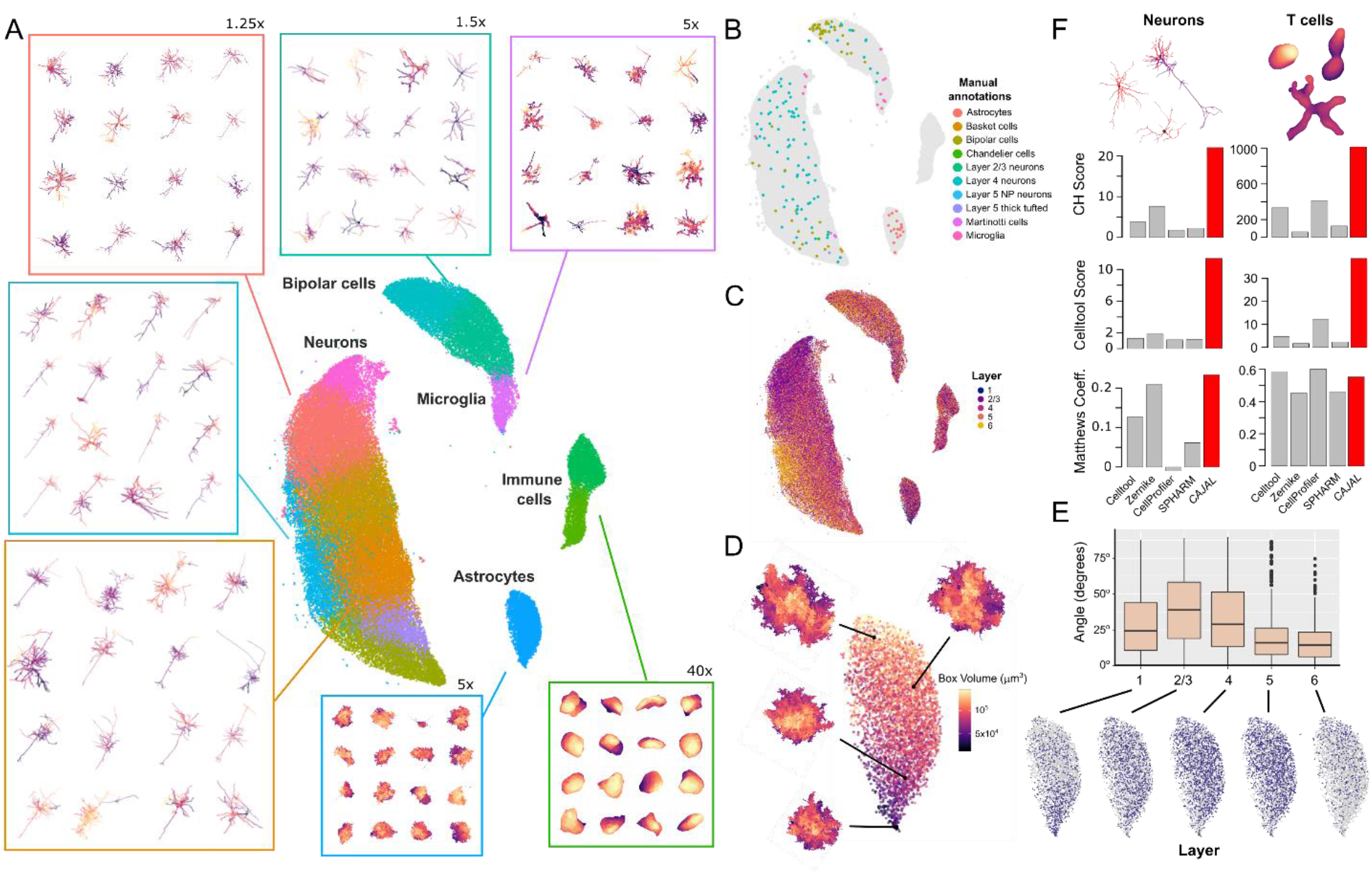
Cell morphology spaces summarize cell shapes across heterogeneous cell types. **A)** UMAP representation of the cell morphology space of 70,510 cells from a cubic millimeter volume of the mouse visual cortex profiled by the MICrONS program using a combination of two-photon microscopy, microtomography, and serial electron microscopy (MICrONS Consortium et al., 2021). The representation is colored by the cell populations identified by clustering of the cell morphology space. The morphology of randomly sampled cells from several populations is also show for reference. The magnification is indicated in cases where the morphology of the cells has been zoomed in to facilitate visualization. The cell morphology space correctly recapitulates the diversity of cell types present in the visual cortex and morphological diversity within each cell type. **B)** The position of 185 cells that were manually annotated by the MICRoNS program is indicated in the UMAP representation, showing consistency with the structure of the cell morphology space. **C)** The UMAP representation of the cell morphology space is colored by the cortical layer of each cell. The morphology space recapitulates morphological differences between neurons and astrocytes from different cortical layers. **D)** The part of the UMAP representation corresponding to astrocytes is colored by the volume of the minimum-size box containing the astrocyte. Astrocytes in the lowe part of the UMAP have smaller dimensions than at the top. For reference, the morphology of 4 astrocytes from different parts of the UMAP is also shown. **E)** Boxplot summarizing the distribution of the angle of the major axis of astrocytes from different layers with respect to the pial surface. Astrocytes from layer 2/3 are elongated perpendicularly with respect to the pial surface. For reference, the part of the UMAP representation corresponding to astrocytes colored by the cortical layer each also shown. **F)** The ability of *CAJAL* to identify morphological differences between neurons of different molecular type and T cells from different anatomical locations is evaluated in comparison to four general algorithms for cell morphometry and according to three different metrics of performance. Contrary to state-of-the-art methods for cell morphometry, the geometric approach implemented in *CAJAL* offered accurate results for both T cells and neurons.

To quantitatively evaluate the ability of GW distance to summarize cell morphologies across different cell types in comparison to existing general approaches for cell morphometry, we considered the 3D morphological reconstructions of 512 T cells from the mouse popliteal lymph node, submandibular salivary gland, and skin, profiled with intra-vital two-photon microscopy (Medyukhina et al., 2020), in addition to the neuronal patch-clamp dataset of the mouse visual cortex (Gouwens et al., 2019). We evaluated the ability of *CAJAL* and 4 other general approaches for cell shape analysis (CellProfiler (McQuin et al., 2018), SPHARM (Brechbühler et al., 1995; Medyukhina et al., 2020), Zernike moments (Khotanzad and Hong, 1990; Pincus and Theriot, 2007), and the PCA-based approach of Celltool (Pincus and Theriot, 2007) and VAMPIRE (Wu et al., 2015)) to predict the anatomical location of each T cell and the Cre driver line of each neuron based on their morphology. In this study, *CAJAL* was again the most capable algorithm at separating the morphologies of T cells from different tissues according to most metrics (Fig. 3F). In addition, all methods for general cell shape analysis except for *CAJAL* performed poorly in the analysis of complex neuronal morphologies, showing that no other method was able to perform well in both datasets (Fig. 3F). Altogether, these results demonstrate that the GW distance overcomes the limitations of current methods to summarize the broad range of complex cell shapes present in mammalian tissues.

### Multimodal analyses of GW cell morphology spaces enable uncovering genetic determinants of cell morphology

The combined analysis of morphological and genomic data from individual cells has the potential to unravel the genetic and molecular pathways that are associated with the progression of high-level cellular processes such as cell differentiation and plasticity. Since changes in cell morphology are continuous, establishing associations between cell morphology and other data is best accomplished by methods of analysis that are purpose-built for continuous processes. We extended our previous work on clustering-independent analyses of omics data (Govek et al., 2019) to implement a statistical approach for identifying molecular and physiological features that are associated with changes in cell morphology. We use the Laplacian score for feature selection (He et al., 2006) to test the association between the values of each feature and the structure of the morphology space, while accounting for user-specified covariates such as the age of the individual (Fig. 4A). To illustrate this approach, we used it to identify genes that contribute to the morphological plasticity of neurons in the *C. elegans*. For that purpose, we considered the 3D morphological reconstructions of the male *C. elegans* GABAergic DVB interneuron in 12 gene mutants, 5 double mutants, and controls across days 1 to 5 of adulthood (Supplementary Table 1), including 7 gene mutants and 1 double mutant from a previous study (Hart and Hobert, 2018). The DVB neuron develops post-embryonically in the dorsorectal ganglion and undergoes post-developmental neurite outgrowth in males, altering its morphology and synaptic connectivity, and contributing to changes in the spicule protraction step of male mating behavior (LeBoeuf and Garcia, 2017). We applied our approach to identify loss of function mutations that are associated with changes in the dynamic morphology of the DVB neuron, taking the age of the worm as a covariate to reliably compare morphological changes across timepoints in adulthood. This analysis identified mutations in the genes *unc-97, lat-2*, *nlg-1, unc-49, nrx-1*, and *unc-25* as significantly affecting the morphology of the DVB neuron (Figs. 4B-D; Laplacian score permutation test, FDR < 0.05). By repeating this analysis for worms of each age separately, we identified the age at which each of these mutations starts significantly affecting the morphology of the DVB neuron (Fig. 4E). To interpret these morphological differences in terms of neuronal characteristics, we used the same approach to evaluate the association of 33 morphological features with the structure of the cell morphology space (Supplementary Table 2). Within these significantly associated features, we found that mutations in *nlg-1* and *unc-25* caused an increase in neurite length and number of branches compared to control worms (Supplementary Fig. 3), while inactivating mutations in *unc-97* and *nrx-1* stunted neurite growth (Supplementary Fig. 3). Altogether, these results are consistent previous findings (Hart and Hobert, 2018), where the morphological phenotype of inactivating mutations if *unc-97*, *nlg-1*, and *nrx-1* was described, and extend them by uncovering new genetic determinants of neuronal plasticity in *C. elegans* and quantitative differences in the age of onset of the morphological alterations induced by different genes.

**Figure 4.**
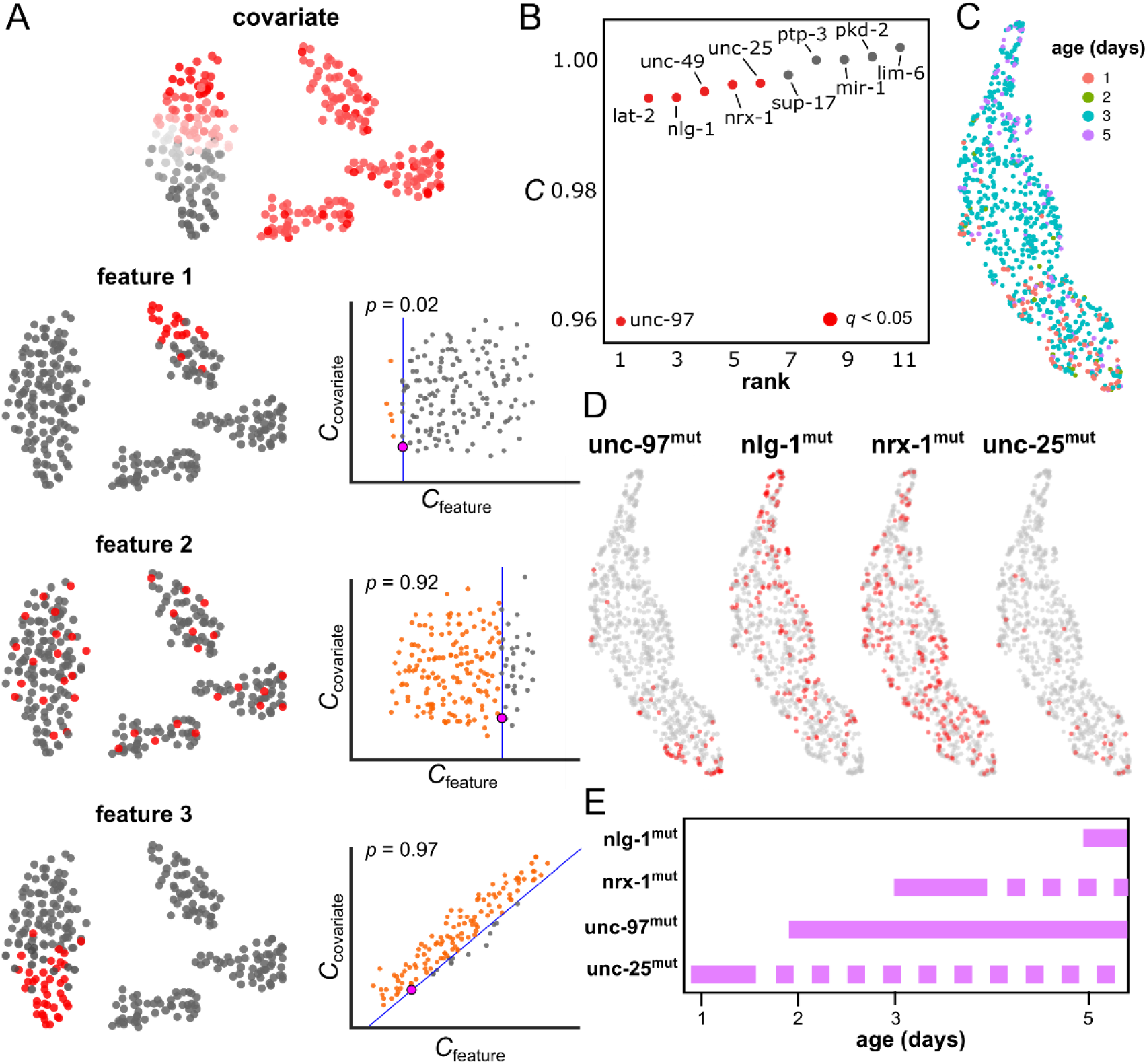
Identification of mutations that have an impact in the morphology of an individual neuron. **A)** Schematic of approach for identifying features (gene expression, mutations, protein expression, etc.) associated with cell morphological changes based on multi- modal data. Features and covariates can take binary (e.g. mutations) or continuous (e.g. gene expression) values. For each feature, the degree of consistency between the feature values and the structure of the cell morphology space is quantified using the Laplacian score (*C*) of the feature in this space. Features with a low score are associated with local regions of the cell morphology space. The statistical significance of each feature in relation to the covariates is evaluated by means of a permutation test, where cell labels are reshuffled. In the figure, examples of features that are significantly localized in the cell morphology space (feature 1, a small number of random configurations have a smaller value of *C*_feature_, independently of the value of *C*_covariate_), not significantly localized in the cell morphology space (feature 2, a large number of random configurations have a smaller value of *C*_feature_), and substantially localized in the morphology space but in association with the covariate (feature 3, a small number of random configurations have smaller value of *C*_feature_, but they are not independent on the value of *C*_covariate_), are presented. **B)** Mutations that have an impact on the morphology of the DVB interneuron in *C. elegans*. Null alleles are ranked according to their Laplacian score (*C*) in the cell morphology space of the DVB interneuron. Samples were imaged across 4 time points and the age of the worm used as a covariate. Genes that significantly impact the morphology of the DVB interneuron according to this approach are indicated in red (FDR < 0.05). **C, D)** UMAP visualization of the cell morphology space of the DVB interneuron colored by the age of each worm (C) and the mutation status of *unc-97, nlg-1*, *nrx-1*, and *unc-25* (D) (red: mutated; gray: wild-type). **E)** Restricting the analysis to worms of the same age allows us to identify the age of onset of the morphological effects induced by each significant mutation (FDR < 0.05). This analysis shows that *unc-97* and *unc-25* mutations have an earlier onset in morphology than *nlg- 1* and *nrx-1* mutations. Dashed lines indicate time points for which there is limited data to restrict the analysis.

### An integrative analysis of molecular, physiological, and morphological data from single cells identifies continuous morpho-transcriptomic trajectories

The incorporation of single-cell RNA-seq onto whole-cell patch-clamp, known as Patch-seq, has enabled concurrent high-throughput measurements of the transcriptome, physiology, and morphology of individual cells (Lipovsek et al., 2021). The integrative analysis of these multi- modal data has the potential to uncover the transcriptomic and physiological programs associated with morphological changes of cells.

We used *CAJAL* to analyze the basal and apical dendrites of 370 inhibitory and 274 excitatory motor cortex neurons profiled with Patch-seq (Scala et al., 2021). Consistent with our previous results, the GW cell morphology space captured morphological differences between the dendrites of neurons from different neuronal t-types and cortical layers (Figs. 5A, B). By representing the pairwise distance between each pair of cells in the transcriptomic, electrophysiological, and morphological latent spaces as a point in a 2D simplex, we found a large degree of variability in the morphology of the dendrites of extratelecephalic-projecting (ET) neurons that was not paralleled by their gene expression profile (Fig. 5C). In contrast, the dendrites of *Lamp5*^+^ and bipolar (*Vip*^+^) GABAergic neurons presented limited morphological variability in comparison to their gene expression profile (Fig. 5C).

**Figure 5.**
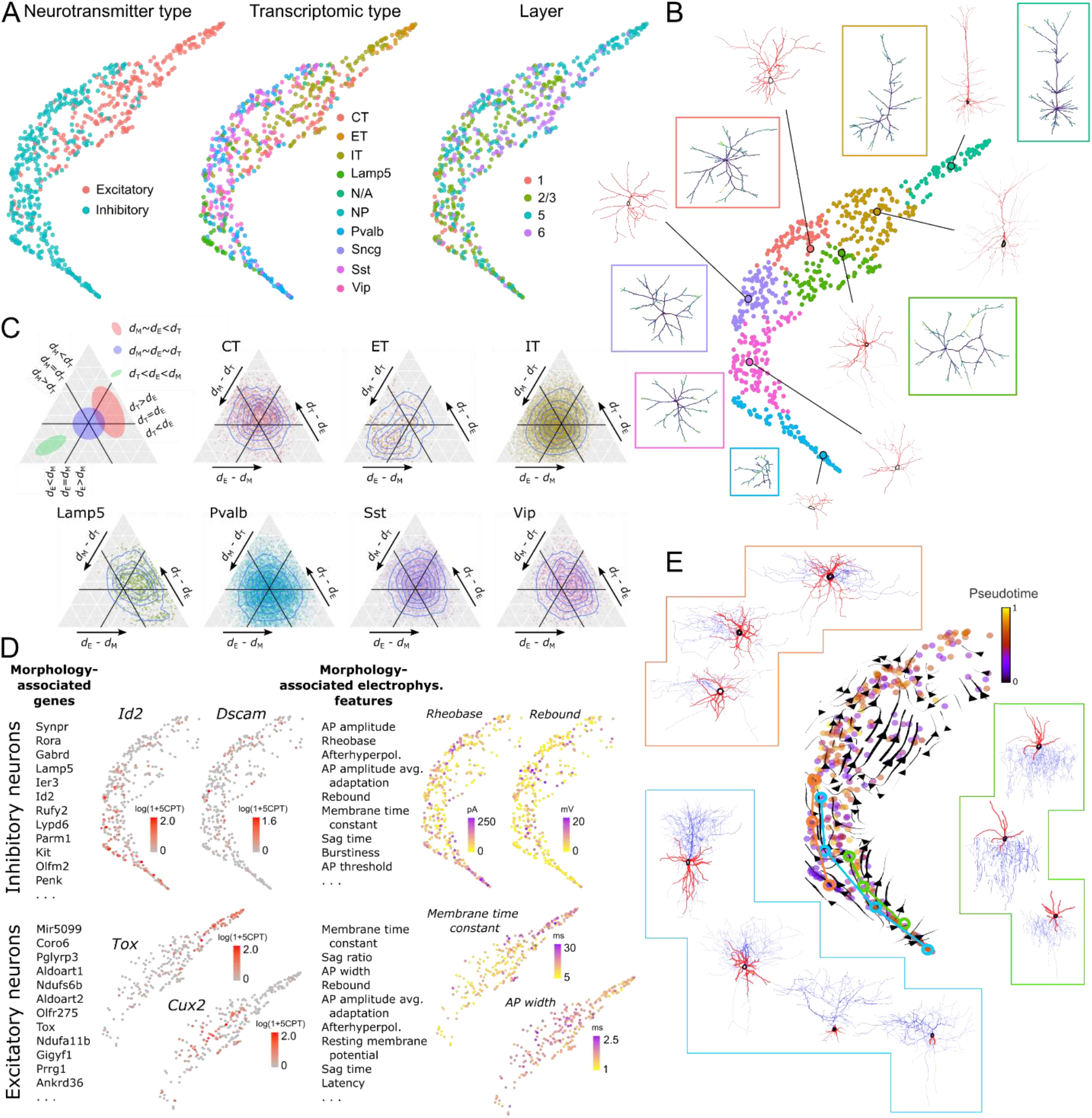
Integrative analysis of molecular, physiological, and morphological data of mouse motor cortex neurons. **A)** UMAP representation of the GW cell morphology space of the dendrites of 370 inhibitory neurons and 274 excitatory neurons from the mouse motor cortex profiled with Patch-seq by (Scala et al., 2021). The representation is colored by the neurotransmitter type (excitatory/inhibitory), the transcriptomic type, and the cortical layer of the cells, showing a large degree of localization of molecular and physiological features on the morphological space. **B)** UMAP representation colored by the morphological cell populations defined by Louvain clustering. The medoid and average cell morphology (in boxes) are shown for each cell population. **C)** Ternary plots showing the discrepancy between pairwise distances between cells in the morphology (M), transcriptomic (T), and electrophysiology (E) latent spaces for each transcriptomically-defined population. The dendrites of ET excitatory neurons present a large degree of variability in their morphology which is not paralleled by consistent changes in their gene expression profile, whereas the dendrites of *Lamp5*^+^ GABAergic neurons present limited morphological variability in comparison to their gene expression profile. CT: corticothalamic neurons, ET: extratelecephalic neurons, IT: intratelenchephalic neurons. **D)** Top genes and electrophysiological features that are significantly associated with the morphological diversity of excitatory and inhibitory neurons according to their Laplacian score in the cell morphology space (FDR < 0.1). The part of the UMAP corresponding to excitatory or inhibitory neurons is colored by the expression level and values of some of the significant genes and electrophysiological features, respectively. CPT: counts per thousand. **E)** Morpho-transcriptomic trajectories computed by projecting the RNA velocity field in the cell morphology space. The morphology of several chandelier, basket, and *Lamp5*^+^ neurons along the trajectories is shown for reference.

We characterized the gene expression and electrophysiological programs associated with morphological differences between neurons by using the Laplacian score approach described above. We performed this analysis separately for inhibitory and excitatory neurons to identify 173 and 556 genes, and 14 and 22 electrophysiological features, respectively, that were significantly associated with the morphological diversity of their dendrites (Fig. 5D and Supplementary Tables 3 and 4; Laplacian score permutation test, FDR < 0.1). Among the 7 genes that were significant for both excitatory and inhibitory neurons, there were several that have been previously reported to be involved in dendrite morphogenesis, such as *Dscam*, which plays a central role in dendritic self-avoidance (Fuerst et al., 2008), and *Pcdh7*, which regulates dendritic spine morphology and synaptic function (Wang et al., 2020). Consistent with these results, a gene ontology enrichment analysis for biological processes identified neuron projection morphogenesis among the top ontologies associated with the morphological diversity of excitatory neurons (GO enrichment adjusted *p*-value = 0.009), and cellular response to chemical stimulus, neuron differentiation, and taxis among the top ontologies associated with the morphological diversity of inhibitory neurons (GO enrichment adjusted *p*-values = 0.006, 0.008, and 0.01, respectively).

We next investigated if some of these gene expression programs form part of continuous morpho-transcriptomic cellular processes. We computed the RNA velocity field to predict the future gene expression state of each cell based on the observed ratio between un-spliced and spliced transcripts (Bergen et al., 2020; La Manno et al., 2018). The time scale of these predictions is determined by the mRNA degradation rate and is of the order of hours. We reasoned that by projecting the RNA velocity field onto the GW cell morphology space and looking for transcriptomic trajectories that also appear as trajectories in this space, we could identify continuous cellular processes that involve consistent changes in gene expression and cell morphology. This approach revealed several morpho-transcriptomic trajectories involving chandelier, basket, and *Lamp5*^+^ neurons (Fig. 5E). Cells along these trajectories showed increased complexity in their apical and basal dendrites in parallel to changes in their gene expression profile, in agreement with the presence of molecular programs associated with the plasticity of these neuronal types. To characterize these molecular programs, we focused our analysis on 78 genes that were associated with the RNA velocity field of inhibitory neurons and computed the Laplacian score of each of these genes in the GW cell morphology space. This analysis revealed that 32 of the 78 genes were also significantly associated with the structure of the cell morphology space (Supplementary Table 5; Laplacian score permutation test, FDR < 0.05). The list of significant genes included multiple genes coding for secreted factors, such as *Spon1*, *Fgf13*, *Rspo2*, and *Reln* (Supplementary Fig. 4), and was enriched for genes involved in memory and cognition (GO enrichment adjusted *p*-value = 0.007).

Taken together, these results demonstrate the utility of *CAJAL* to identify and characterize molecular and electrophysiological programs associated with cell morphological changes based on single-cell Patch-seq data.

### GW cell morphology spaces enable the integration of cell morphology data across technologies

Advances in cell morphology profiling techniques have led to an explosion of high-resolution cell morphology data over the past decade (Ascoli et al., 2007). The ability to perform integrated analyses of such data regardless of the experimental approach and technology that was used to generate them would be a powerful tool for imputing missing data and refining taxonomic classifications of cells. For example, by integrating patch-clamp and Patch-seq data the transcriptome of cells profiled with patch-clamp could be predicted based on their morphology.

We used *CAJAL* to build a combined morphology space of the basal and apical dendrites of visual cortex neurons profiled with patch-clamp (Gouwens et al., 2019) and visual and motor cortex neurons profiled with Patch-seq (Gouwens et al., 2020; Scala et al., 2021). The combined dataset consisted of 1,662 neurons, of which 1,156 had associated single-cell RNA-seq data. Inhibitory and excitatory neurons from different datasets clustered together in separated regions of the combined morphology space (Fig. 6A), indicating that the structure of this space is mostly driven by biological differences rather than by technical differences. To evaluate the consistency of the combined cell morphology space, we considered the t-type (Tasic et al., 2018) of the cells profiled with Patch-seq, and quantitatively assessed the distance in the combined cell morphology space between cells of the same t-type (for the Patch-seq data) or cells labelled with the corresponding Cre driver line (for the patch-clamp data). Cells of same t-type but from different Patch-seq datasets, as well as cells from the matching Cre driver line in the patch-clamp dataset, were closer to each other in the combined morphology space than cells from different t-types or Cre driver lines (Fig. 6B and Supplementary Fig. 5; Wilcoxon rank-sum test *p*-value < 10^-100^), demonstrating the utility of the GW distance for integrating cell morphology data across experiments and technologies.

**Figure 6.**
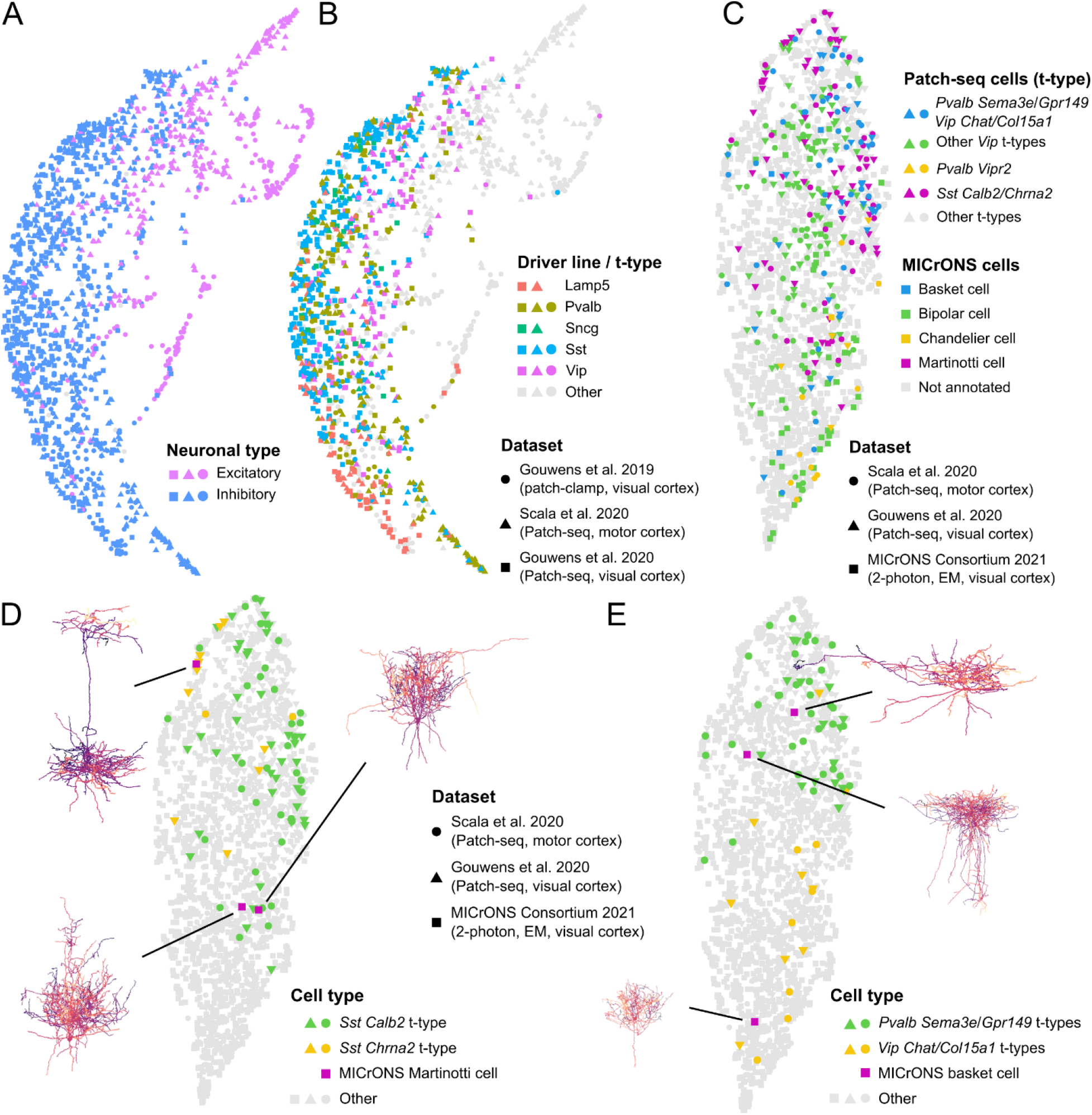
Integration of cell morphology data across experiments and technologies. **A)** UMAP representation of the combined cell morphology space of the basal and apical dendrites of visual cortex neurons profiled with patch-clamp (Gouwens et al., 2019), and visual cortex and motor cortex neurons profiled with Patch-seq (Gouwens et al., 2020; Scala et al., 2021). The combined dataset consists of 1,662 neurons. The representation is colored by the neuronal type. Inhibitory and excitatory neurons from different datasets cluster together in separated regions of the combined morphology space. **B)** The same UMAP representation is colored by the Cre driver line (for cells from the patch-clamp dataset) or the t-type (for cells from the Patch-seq datasets). Cells of same t-type but from different Patch-seq dataset, and cells from the corresponding Cre driver line in the patch-clamp dataset, localize in the same regions of the combined morphology space. **C)** UMAP representation of the combined cell morphology space of 883 full neuron reconstructions from the motor and visual cortices profiled with Patch-seq (Gouwens et al., 2020; Scala et al., 2021) and 1,139 neurons from the mouse visual cortex with a combination of two-photon microscopy, microtomography, and serial electron microscopy (MICrONS Consortium et al., 2021), 139 of which have been manually annotated by the MICrONS program. The manually annotated neuronal types from the MICrONS dataset localize in the same regions of the morphology space than Patch-seq cells from the corresponding t-type. **D)** Refined annotation of 3 Martinotti cells that were manually annotated by the MICrONS program. One of the Martinotti cells presents a distinct morphology with a densely arborized axon and is close in the cell morphology space to Patch-seq cells of the *Sst Chrna2* t-type, while the other two Martinotti cells are closer to *Sst Calb2* t-type cells. **E)** Refined annotation of 3 basket cells that were manually annotated by the MICrONS program. One basket cell has a more condensed morphology than the others is close in the morphology space to Path-seq cells of the *Vip Chat Htr1f* and *Vip Col15a1 Pde1a* t-types, while the other two larger basket cells are close to *Pvalb Sema3e Kank4* and *Pvalb Gpr149 Islr* t-type cells.

We then used the same approach to refine the annotation of visual cortical neurons profiled with serial electron microscopy by the MICrONS program (MICrONS Consortium et al., 2021). We considered 883 full neuron reconstructions from the two Patch-seq datasets and created a combined morphology space of these cells along with a subset of 1,000 evenly sampled and 139 manually annotated neurons from the MICrONS dataset. As with the integration of patch-clamp and Patch-seq data, the manually annotated neuronal types from the MICrONS dataset were closer to Patch-seq cells of the matching t-type in the consolidated cell morphology space than to non-matching t-types (Fig. 6C and Supplementary Fig. 6; Wilcoxon rank-sum test *p*- value < 10^-100^). For example, the only chandelier cell annotated in our MICrONS dataset was closer to *Pvalb Vipr2* t-type cells from the Patch-seq data than to cells from other t-types (Supplementary Fig. 6; Wilcoxon rank-sum test *p*-value = 0.035).

Using this combined cell morphology space, we refined some of the manual annotations of the MICrONS data with more precise transcriptomic definitions. For example, of the three Martinotti cells annotated in the MICrONS dataset, one cell presenting a distinct morphology with a densely arborized axon was closer in the morphology space to Patch-seq cells of the *Sst Chrna2* t-type (Fig. 6D; Wilcoxon rank-sum test *p*-value = 0.045), while the other two Martinotti cells were closer to *Sst Calb2* t-type cells (Fig. 6D; Wilcoxon rank-sum *p*-value = 10^-3^). This is consistent with previous results showing that expression of *Chrna2* is characteristic of layer 5 Martinotti cells that project into layer 1 (Hilscher et al., 2017), and we confirmed that the soma of the predicted *Chrna2* Martinotti cell was indeed located in layer 5 while its long axon ended in layer 1 (Fig. 6D). Similarly, among the manually annotated basket cells in the MICrONS dataset, one had a more condensed morphology than the others (Fig. 6E). This smaller basket cell was close in the cell morphology space to *Vip Chat Htr1f* and *Vip Col15a1 Pde1a* t-type Patch-seq cells (Fig. 6E; Wilcoxon rank-sum *p*-value = 0.02), while larger basket cells were closer to *Pvalb Sema3e Kank4* and *Pvalb Gpr149 Islr* t-type Patch-seq cells (Fig. 6E; Wilcoxon rank-sum *p*- value = 10^-14^). These results were again in agreement with the molecular characterization of small and large basket cells in the somatosensory cortex (Wang et al., 2002).

Taken together, these results demonstrate the utility of GW cell morphology spaces to perform integrative analyses of cell morphological data across technologies and represent a conceptual basis for the development of algorithms for cell morphological data integration.

## Discussion

Shape registration has experienced several breakthroughs over the past 15 years with the formalization of new paradigms that allow for more flexibility in the quantification of morphology (Biasotti et al., 2016). Here, we built upon one of these constructions, the GW distance, to develop a general computational framework and software for the multi-modal analysis and integration of single-cell morphological data. The proposed framework does not rely on predefined morphological features, is insensitive to rigid transformations, and can be efficiently used with arbitrarily complex and heterogeneous cell morphologies. Using this approach, we have accurately built, analyzed, and visualized cell morphology latent spaces. Similar to gene expression latent spaces in the analysis of single-cell RNA-seq data, morphological latent spaces are instrumental in the analysis and integration of single-cell morphological data. The metric properties of these spaces allowed us to integrate single-cell morphological data of neurons across experiments and technologies; identify morphological, molecular, and physiological features that define different subpopulations of neurons and glia; and establish associations between morphological, molecular, and electrophysiological cellular processes in these cells. Our quantitative and comparative studies using Patch-seq, patch-clamp, electron, and two-photon microscopy data demonstrate that GW-based morphological analyses represent a substantial improvement in accuracy and scope with respect to current methods for the analysis of cell morphology data, and enable previously unavailable analyses, such as inferring the transcriptional type of individual neurons based on the morphology of their dendrites.

More generally, we expect that the analytic framework presented in this work will serve as the basis for the development of other currently missing computational methods for the analysis of single-cell morphological data, such as methods for batch-correcting cell morphological data or modelling morphological processes. The development of these and other methods for single-cell morphological data analysis will significantly impact our understanding of the relation between the morphological, molecular, and physiological diversification of cells.

## Methods

### Computation of GW cell morphology spaces

We build upon the application of metric geometry to the problem of finding a correspondence between two point-clouds such that the size of non-isometric local transformations is minimized (Mémoli, 2007, 2011; Mémoli and Sapiro, 2005). *CAJAL* takes as input the digitally reconstructed cells. For each cell *i*, it samples *n* points regularly from the outline and computes their pairwise distance matrix, *d_i_*. It then computes the GW distance between every pair of distance matrices

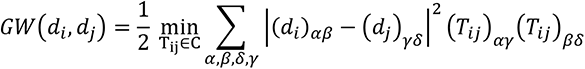

where the matrix *T_ij_* specifies a weighted pointwise matching between the points of cells *i* and *j*, and *C* represents the space of all possible weighted assignments (Mémoli, 2011). By construction, *CAJAL* does not require pre-aligning cell outlines, since the input to *GW* is the pairwise distance matrix within each cell, *d_i_*, which is invariant under rigid transformations. Depending on the application, we consider two choices for the distances *d_i_*: Euclidean and geodesic distance.

The output is a metric space for cell morphologies which can then be clustered and visualized using standard procedures, such as Louvain community detection (Blondel et al., 2008) and UMAP (Becht et al., 2018). For each population of cells, *x*, we compute its average morphology as the distance matrix

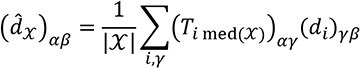

where med(*x*) denotes the medoid of *x* with respect to the *GW* distance matrix. The morphology can then be visualized by computing the shortest-path tree or multidimensional scaling (MDS) of *d_x_*. In addition, to facilitate the interpretation of morphology spaces, we find it is useful to plot the values of standard morphological descriptors like cell height, width, diameter, neuronal depth, or fractal dimension (Arshadi et al., 2021; McQuin et al., 2018; Scorcioni et al., 2008) in the UMAP representation of the cell morphology space.

### Evaluation of features on the cell morphology space

To evaluate features, such as gene expression or electrophysiological properties, on GW cell morphology spaces, we build upon a spectral approach for clustering-independent analyses of multimodal data (Govek et al., 2019; He et al., 2006). We first construct a radius neighbor graph of the *GW* distance with radius ε. Each feature *g* is represented by a vector *f_g_* of length the number of cells.

The Laplacian score of *g* on the cell morphology space is then given by (He et al., 2006)

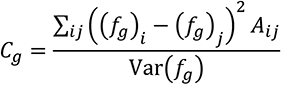

where *A* is the adjacency matrix of the radius neighbor graph, and Var(*f_g_*) the estimated variance of *f_g_*. Features with a low *C_g_* score are associated with morphologically similar cells. The significance of the score can be statistically assessed for each feature by means of a one-tailed permutation test and adjusted for multiple hypothesis testing using Benjamini-Hochberg procedure. To assess the significance of a feature *g* in the presence of a set of covariates *h_m_*, we perform a permutation test where the entries of *f_g_* and *f_h_m__* are simultaneously permuted and the scores *C_g_* and *C_h_m__* are computed at each permutation. We denote these values collectively as 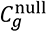 and 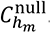. We then solve the regression problem

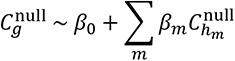

and consider the distribution of residuals as the null distribution for the adjusted score 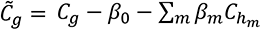.

### Processing of Patch-seq and patch-clamp morphological reconstructions

We downloaded the morphological reconstructions of neurons from several repositories. For the Patch-clamp dataset of (Gouwens et al., 2019), we downloaded 509 reconstructions in SWC format from the Allen Cell Types database, using Cell Feature Search and selecting for “Full” or “Dendrite Only” reconstruction types. Three of the SWC files were unsorted and were left out of further processing, for a total of 506 neurons. For the Patch-seq dataset of (Gouwens et al., 2020), we downloaded 574 reconstructions from the Brain Image Library (BIL) repository. We removed 62 neurons that did not have assigned transcriptomic types, for a total of 512 neurons. Lastly, for the Patch-seq dataset of (Scala et al., 2021), we downloaded 645 reconstructions from the inhibitory and excitatory sets from the BIL repository, skipping the one inhibitory neuron that had no dendrites, for a total of 644 neurons.

The SWC format represents each neuron as a tree of vertices, such that an edge can be drawn between a vertex and its parent, forming the skeleton of the neuron. From this format, we sampled 100 points radially around the soma at a given step size. We used a binary search to identify the step size which returns the required amount of evenly spaced points. To calculate the pairwise geodesic distance between these points, we constructed a weighted graph with weights given by the distance to the latest sampled point. We then used the Floyd-Warshall algorithm implemented in Networkx (Hagberg et al., 2008) to compute all pairwise shortest path distances in this graph. Alternatively, we computed the pairwise Euclidean distance between the 3D coordinates of these points.

We the computed the GW distance between each pair of cells as described above (subheading “Computation of GW cell morphology spaces”) using the ot.gromov.gromov_wasserstein function of the “POT: Python Optimal Transport” Python library (Flamary et al., 2021). We then used this precomputed distance to build a 2D visualization of the morphology space using the https://github.com/tkonopka/umap package in R. We computed the Louvain clusters of a KNN graph of the GW distance using the multilevel.community function of the igraph R package (Csardi and Nepusz, 2006).

### Average shape of neurons

To compute the average shape of a cluster of cells in the GW cell morphology space, we first found the medoid cell as the cell with the minimum sum of distances to all other cells in the cluster. To compute a morphological distance between cells, the GW algorithm identifies an optimal matching *T_ij_* between the points we have sampled (subheading “Computation of GW cell morphology spaces”). We used this matching to align the other cells in the cluster to the medoid, by reordering the pairwise geodesic distance matrix of their sampled points to match the distance matrix of the medoid cell. We rescaled the geodesic distance matrix of each cell into an unweighted graph distance by dividing out the minimum distance between any two points, so that the rescaled distances were integers. We set a threshold on these distances at 2, such that the distance was 0 from the point to itself, 1 to an adjacent point in the tree of the neuron trace, and 2 to any farther point. We averaged all of these distance matrices together over the cells in the cluster and built a *k* = 3 nearest neighbor graph, essentially connecting each sampled point to the three other points it was most often adjacent in the neurons of that cluster. We took the shortest path tree in this graph as the average shape for that cluster using Dijkstra’s algorithm. We color each point in this average shape by a confidence value based on its minimum original unweighted graph distance, summed over the cluster, to any other point.

### Comparison of *CAJAL* with current methods for neuronal morphometry

We compared our approach to five other morphological methods for neuron analysis by applying them to the dendrite reconstructions of the neuronal datasets listed above (subheading “Processing of Patch-seq and patch-clamp morphological reconstructions”). These methods have stricter assumptions on the input, forcing us to remove disconnected dendrites from the reconstructions. We applied NBLAST (Costa et al., 2016), as implemented in the nat.nblast R package (https://github.com/natverse/nat.nblast). We calculated a pairwise distance between all neurons using the nblast_allbyall function with the mean normalization method. We ran the Topological Morphology Descriptor (TMD) method of (Kanari et al., 2018) using the TMD Python package (https://github.com/BlueBrain/TMD). We followed their distances example (https://github.com/BlueBrain/TMD/blob/master/examples/distances_example.py) to compute the persistence image difference between every pair of neurons. We skipped 26 neurons across the two Patch-seq datasets for which get_ph_neuron or get_persistence_image_data errored due to a lack of bifurcating branches. We used the Measure Multiple Files batch script of the ImageJ SNT plugin (Arshadi et al., 2021) to compute morphological features of neurons, including the Sholl features (Sholl, 1953). We also computed morphological features using L-Measure (Scorcioni et al., 2008), selecting all of their provided functions.

We used three different metrics to assess the ability of these algorithms to identify morphological differences between Cre lines or transcriptomic types. We computed the Calinski-Harabasz clustering score using cluster.stats from the fpc R package (https://cran.r-project.org/package=fpc). We also implemented the median-based group discrimination statistic used by (Pincus and Theriot, 2007) to compare methods for cell-shape analysis. Lastly, we used a 7-fold *k* = 10 nearest neighbor classifier from the scikit-learn Python library to predict the Cre driver line or t-type of each cell based on morphological distance and used the Matthew’s correlation coefficient to evaluate the accuracy of the predictions.

### Morphological analysis of the MICrONS dataset

We downloaded the 113,182 static cell segmentation meshes from MICrONS using the trimesh_io module from the package MeshParty (https://meshparty.readthedocs.io/) at the lowest resolution (resolution 3). We then downloaded higher resolution meshes for cells that had less than 10,000 vertices at this lowest resolution. Cells with less than 1,001 vertices at the lowest resolution were re-downloaded at the highest resolution (resolution 0). Cells with 1,001 to 3,000 vertices at the lowest resolution were re-downloaded at resolution 1, and cells with 3,001 to 10,000 vertices were re-downloaded at resolution 2.

Along with other metadata available through the CAVEclient (https://github.com/seung-lab/CAVEclient), such as the 3D coordinates of neuron soma, we collected the cell IDs for each manually annotated cell type provided by the MICrONS program in their website. We used the layer 2/3, layer 4, and layer 5 manually annotated cells to estimate cortical layer boundaries in the y values of the 3D soma coordinates. We placed these cutoffs at layer 1 < 104,191 < layer 2/3 < 133,616 < layer 4 < 179,168 < layer 5 < 213,824 < layer 6.

We sampled 50 vertices from the triangular mesh of each cell, using the linspace function of the NumPy package (Harris et al., 2020) to evenly select vertices, since vertices were roughly ordered by proximity, and this gives an approximation of even sampling over the 3D space. We skipped the very large blood vessel mesh and 240 meshes with less than 50 vertices, for a total of 112,941 meshes. We used the heat method (Crane et al., 2013), implemented in the MeshHeatMethodDistanceSolver function of the Python library potpourri3D (https://github.com/nmwsharp/potpourri3d), to compute geodesic distances between the sampled points on the mesh. We parallelized the computation of the pairwise GW distance between the 112,941 meshes on 128 cores, but otherwise used the same process as with the Patch-seq and patch-clamp datasets (subheading “Processing of Patch-seq and patch-clamp morphological reconstructions”). Due to the large size of the resulting GW pairwise distance matrix, we used the Python libraries leidenalg (Traag et al., 2019) and umap-learn (Leland et al., 2018) to cluster the cells and compute 2D UMAP visualizations, respectively. We labelled the clusters based on the manually annotated cells provided by the MICrONS program.

We found that some morphological clusters mostly consisted of artifacts or neuron-glia doublets and removed those. In addition, another morphological cluster contained both neuron-neuron doublets and individual neurons with complex morphologies, so we devised an approach to remove meshes containing multiple somas from that cluster. We determined the number of somas in each mesh from that cluster by using MeshParty to skeletonize the meshes and convert them into graph representations where nodes have a radius value, and nodes within soma regions fall in a specific range of radii. For us, this range was 4,000 to 30,000. We used HDBSCAN (McInnes et al., 2017) to cluster these nodes in the 3D space and counted each cluster with at least three nodes as a soma. Meshes with more than one soma were removed from the cluster. Lastly, we noticed that many meshes with very high y coordinates appeared stretched, so we removed meshes with a y soma coordinate greater than 240,000. After removing all these artifacts, we recomputed a UMAP visualization of the remaining 70,510 cells in the cell morphology space using umap-learn.

For each astrocyte, we measured the bounding box by placing lower and upper bounds on the 1% and 99% quantiles of the mesh vertices along each of the first three principal components. We took the arccosine of the first principal component along the y axis to be the orientation angle of the astrocyte and measured its deviation from perpendicular.

### Morphological analysis of T cells

We retrieved 512 3D meshes of T cells from (Medyukhina et al., 2020). We evenly sampled 200 points from the list of vertices in each mesh, which approximates an even sampling in 3D space since the vertices are roughly ordered in a spiral down the cell. We computed the GW distance of the pairwise Euclidean distances between these points as described above (subheadings “Computation of GW cell morphology spaces” and “Processing of Patch-seq and patch-clamp morphological reconstructions”).

### Comparison of CAJAL with general methods for cell morphometry

We applied the Celltool method of (Pincus and Theriot, 2007) using their Python package (https://github.com/zpincus/celltool). Since this method only works with 2D cell segmentations, we sampled the 2D boundary of the projection of each cell along the first two axes to the same number of sampled points used for *CAJAL*. We aligned these contours using a maximum of 20 iterations, allowing for reflections, and saved the non-normalized PCA values from the shape model. We used CellProfiler 4.0.3 (Carpenter et al., 2006) on binary 2D projection images to compute both general shape features and Zernike moments using MeasureObjectSizeShape. We ran SPHARM (Medyukhina et al., 2020) using their Python package (https://github.com/applied-systems-biology/Dynamic_SPHARM) on all of the mesh vertices for each cell. For neurons, we used the marching_cubes function of the scikit-learn Python library to define 3D mesh vertices. We used the spectrum.return_feature_vector function of SPHARM to extract the amplitude of harmonic components from the spectra produced by compute_spharm. We compared these methods to *CAJAL* using the same metrics described above (subheading “Comparison of *CAJAL* with current methods for neuronal morphometry”).

### Evaluation of the accuracy and runtime of *CAJAL* as a function of the number of sampled points

We sampled 25, 50, 75, 100, and 200 points from each cell from the patch-clamp dataset of (Gouwens et al., 2019) and applied *CAJAL* as described above to compute the GW distance between cells. We used the Calinski-Harabasz score, the median-based statistic of (Pincus and Theriot, 2007), and the Matthews coefficient of a *k* = 10 nearest neighbor classifier (subheading “Comparison of *CAJAL* with current methods for neuronal morphometry”) to assess how the number of sampled points affects the ability of *CAJAL* to capture morphological differences between cells from different Cre driver lines. Runtimes were determined based on 12 threads of a desktop computer with an 8-core Intel Xeon E5-1660 3.20 GHz CPU.

### Morphological analysis of the DVB neuron

We considered the neurite reconstructions of the DVB neuron from 799 adult male *C. elegans* aged 1 to 5 days from control strains or strains containing mutations in the genes *nrx-1, mir-1, unc-49, nlg-1, unc-25, unc-97, lim-6, lat-2, ptp-3, sup-17,* or *pkd-2* (Supplementary Table 2). Reconstructions were created from confocal images of the DVB neuron using SNT (Arshadi et al., 2021) in Fiji (Schindelin et al., 2012) as previously described (Hart and Hobert, 2018). We computed the GW distance between these morphological reconstructions as described above (subheadings “Computation of GW cell morphology spaces” and “Processing of Patch-seq and patch-clamp morphological reconstructions”), based on the geodesic pairwise distance of 100 points sampled from each neuron. We then introduced an indicator function for each mutated gene, which took values 0 or 1 on each cell depending on whether the worm had a wild-type or a mutated version of the gene, respectively. To determine which of the 11 mutated genes were associated with changes in morphology, we computed the Laplacian score of each indicator function on the GW cell morphology space as described above (subheading “Evaluation of features on the cell morphology space”). To compute the score we used the R package RayleighSelection (Govek et al., 2019) with 1,000 permutations, *ε* equal to the median GW distance and the age of the worm in days as a covariate. In the same way, we used RayleighSelection to determine which of 33 morphological features computed with SNT were significantly localized in the cell morphology space. In addition, we performed the same analysis using only neurons from a single day, for each day, to determine the age at which the effect of significant mutations on the morphology of the DVB neuron starts to emerge.

### Identification of genes and electrophysiological features associated with the morphology of neuronal dendrites

We used the same process described above (subheading “Processing of Patch-seq and patch-clamp morphological reconstructions”) to compute the GW distance between the morphological reconstructions of the dendrites of 644 neurons profiled by (Scala et al., 2021). We sampled 100 points from each dendrite and used geodesic distance to measure the distance between points. To determine which genes are associated with morphological variability we computed the Laplacian score of each gene on the GW morphology space using RayleighSelection, as described above (subheading “Evaluation of features on the cell morphology space”). Gene expression values were normalized as log(1 + 5000 ·size-normalized expression), we used 1,000 permutations, and ε was given by the median GW distance. We only tested genes expressed in at least 5% and less than 90% of cells. We identified gene ontology enrichments using the R package gProfileR (Reimand et al., 2007), where we performed an ordered query of the significant genes based on their Laplacian score and restricted the search to biological process (BP) gene ontologies. We used the same procedure based on the Laplacian score to determine which electrophysiological features were associated with changes in the morphology of the dendrites.

### Computation of RNA velocity trajectories

We clipped 3’ Illumina adapters and aligned FASTQ files to the GRCm38 mouse reference genome using the STAR aligner (Dobin et al., 2013). We used the command “run_smartseq” from the velocyto command line tool (La Manno et al., 2018) to create a Loom file of spliced and unspliced reads. We then used the scvelo Python package (Bergen et al., 2020) to compute RNA velocity trajectories. We tested scvelo in dynamical or stochastic mode with 0, 10, or 20 minimum counts; 500, 1000, or 2000 top variable genes; 10, 20, or 30 principal components; and 10 or 30 neighbors. We kept the velocity trajectories with the highest average confidence per arrow, defined by agreement with neighboring arrows. These trajectories were produced using stochastic mode with 0 minimum counts, 500 top variable genes, 10 principal components, and 30 neighbors. We computed the pseudotime using the velocity graph. We took all 78 genes which passed the basic default filters in rank_velocity_genes() to be velocity-related genes and used the Laplacian score to assess their morphological association.

### Consistency between transcriptomic, electrophysiological, and morphological spaces

We defined the transcriptomic distance (*d_T_*) between two cells as the Spearman correlation distance between their log-normalized gene expression profile, and their electrophysiological distance (*d_E_*) as the Euclidean distance between their electrophysiological feature vectors. We compared these distances and the GW morphological distance (*d_M_*) between all pairs of cells in the dataset of (Scala et al., 2021) by representing them on a 2-simplex. For that purpose, we standardized the logarithm of pairwise distances independently for each data modality. We then took the axes of the 2-simplex to be the given by the difference between each pair of distances (*d_M_* – *d_T_*, *d_T_* – *d_E_*, *d_E_* – *d_M_*), so that the sum of the coordinates equals 0 for each pair of cells. We plotted cell pairs in the middle 98% of each axis.

### Integrative analysis of Patch-seq and patch-clamp data

We combined the patch-clamp and two Patch-seq datasets into one cell morphology space by computing the GW distance between the morphological reconstructions of the dendrites of all 1,662 neurons from the 3 datasets. We sampled 100 points from each dendrite and used geodesic distance to measure distances between points.

To evaluate the integration of the Patch-seq datasets, we utilized the classification of neurons into the t-types of (Tasic et al., 2018). This classification is provided by (Gouwens et al., 2020) as their transcriptomic alias, and we computed the classification for the dataset of (Scala et al., 2021) using their ttype-assignment Jupyter notebook. We tested the overlap between neurons of the same t-type but from different datasets in the cell morphology space by performing a Wilcoxon rank-sum test, comparing the distribution of GW distances within the same t-type with the distribution of GW distances between t-types.

To evaluate the integration between the two Patch-seq datasets and the patch-clamp dataset, we matched the neuronal t-types in the Patch-seq datasets with the Cre driver lines in the patch-clamp dataset. We used only the first marker in the t-types and considered markers that existed in at least five cells of two of the three datasets. This left Sst, Pvalb, and Vip as major markers between the t-types and Cre lines, and Lamp5 and Sncg as markers between t-types only. We again used the Wilcoxon rank-sum test to compare the distributions of GW distances within and between these five major transcriptomic types.

### Integrative analysis of Patch-seq and MICrONS neuronal data

We calculated a combined GW morphological space for the two Patch-seq datasets and 1,000 neurons evenly sampled from the MICrONS dataset, in addition to 140 manually annotated neurons by the MICrONS program. We sampled 50 points from the full neuronal reconstructions from the Patch-seq datasets. In the case of the dataset of (Scala et al., 2021), this restricted our analysis to 370 neurons with full reconstructions. Since the SWC format used in the Patch-seq datasets contains a trace reconstruction, and the triangular cell segmentation meshes used in the MICrONS dataset contain cell surface reconstructions, we computed the GW distance based on the pairwise Euclidean distances between 50 points sampled from each neuron, instead of geodesic distance.

To evaluate the integration, we matched some of the manually annotated cells from the MICrONS dataset with t-types from the Patch-seq datasets. Following the results of Tasic *et al*. (Tasic et al., 2018), we assigned the *Sst Calb2/Chrna2* t-types (*Sst Calb2 Pdlim5, Sst Calb2 Necab1, Sst Chrna2 Ptgdr, Sst Chrna2 Glra3*) to Martinotti cells, and the *Pvalb Vipr2* t-type to chandelier cells. Some other *Pvalb* t-types were assigned to basket cells, such as *Pvalb Sema3e Kank4* and *Pvalb Gpr149 Islr,* whereas CCK or small basket cells were associated with *Vip* t-types such as *Vip Chat Htr1f* and *Vip Col15a1* (Wang et al., 2002). Since cells of the *Vip* subclass have bipolar morphologies (Tasic et al., 2018), we assigned all other *Vip* subtypes to bipolar cells. We then evaluated the consistency of the cell morphology space with these assignments by using a Wilcoxon rank-sum test to compare the distribution of GW distances between matching types across datasets with the distribution GW distances between non-matching types across datasets.

### Code availability

The source code of *CAJAL* is available at https://github.com/CamaraLab/CAJAL.

### Data availability

All the datasets used in this study are publicly available. The morphological reconstructions of the DVB neuron have been deposited in the neuromorpho.org database (Hart archive). The patch-clamp data of (Gouwens et al., 2019) can be accessed at the Allen Brain Atlas data portal (http://celltypes.brain-map.org/data). The Patch-seq datasets of (Gouwens et al., 2020) and (Scala et al., 2021) can be accessed at the Brain Image Library (BIL) using the URLs https://download.brainimagelibrary.org/biccn/zeng/pseq/morph/200526/ and https://download.brainimagelibrary.org/biccn/zeng/tolias/pseq/morph/, respectively. The two-photon microscopy data of (Medyukhina et al., 2020) can be accessed at https://asbdata.hki-jena.de/publidata/MedyukhinaEtAL_SPHARM/. The MICrONS program dataset can be accessed using the MICrONS Explorer (https://www.microns-explorer.org/cortical-mm3#segmentation-meshes).

## Acknowledgements

The authors are grateful to Dr. Zhaolan Zhou for his constructive comments on the manuscript, Matthew Jozwik for assisting with the conversion of files for the DVB analysis, and Jiazhen Rong and Alice Wang for collaboration on related aspects.

## Author contributions

K.G. implemented *CAJAL* and performed all the computational analyses of this work. J.C. performed preliminary computational analyses for this work. A.B.S. implemented the covariate analysis of the Laplacian score. K.Z. and M.P.H. generated the morphological reconstructions of the DVB neuron and assisted with their analysis. P.G.C. and K.G. jointly wrote the manuscript. P.G.C. supervised the work.

