## Supplementary Figures for "Multi-modal analysis and integration of single-cell morphological data"

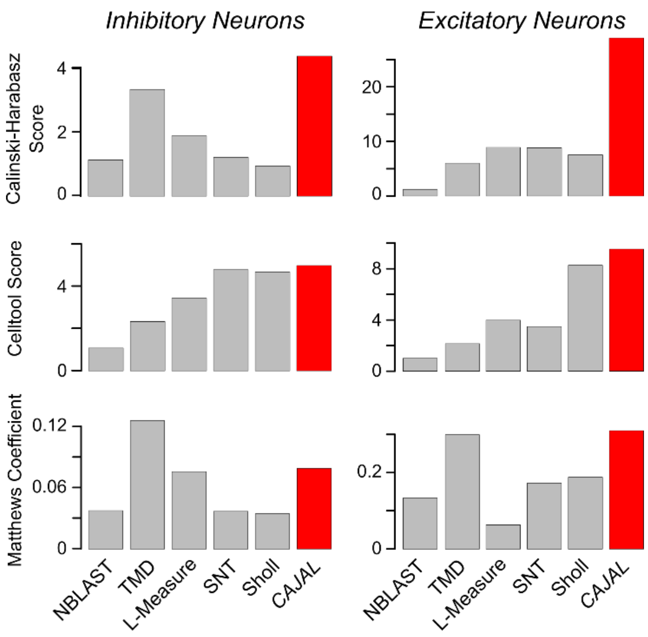


**Supplementary Figure 1. Prediction of excitatory and inhibitory neuron t-types based on the morphology of the morphology of apical and basal dendrites.** The ability of *CAJAL* to identify morphological differences between neurons of different t-types is assessed in 370 inhibitory and 274 excitatory motor cortical neurons (spanning 47 and 26 t-types, respectively) profiled with Patch-seq (Scala et al., 2021) in comparison to 5 state-of-the-art methods for neuronal morphometry.

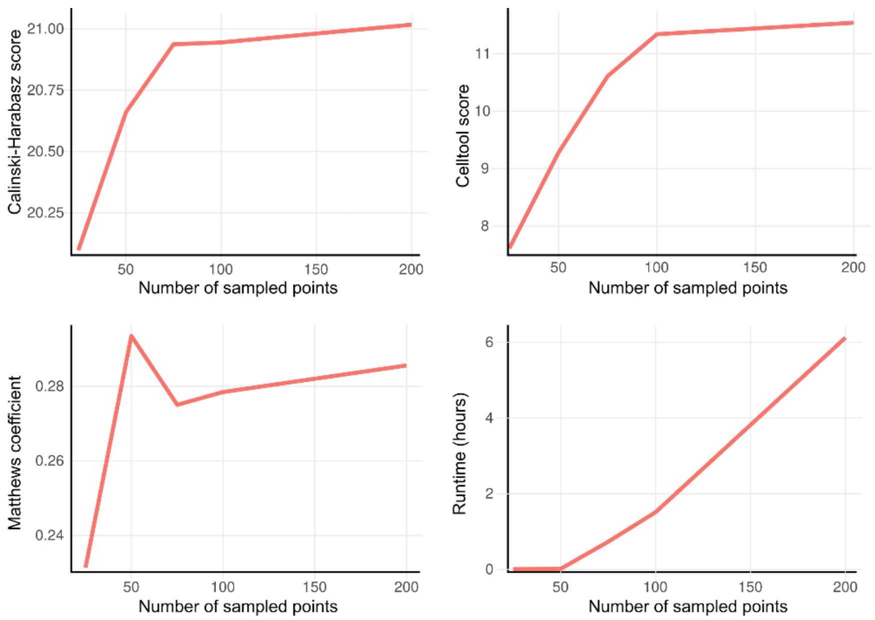


**Supplementary Figure 2. Accuracy and runtime of *CAJAL* as a function of the number of sampled points per cell.** The Calinski-Harabasz score, the Celltool score (Pincus and Theriot, 2007), and the Matthews coefficient of a $k = 10$ nearest neighbor classifier were used to evaluate the ability of *CAJAL* to capture morphological differences between visual cortex neurons labelled with different Cre driver lines (Gouwens et al., 2019). Runtime was determined based on 12 threads of a desktop computer with an 8-core Intel Xeon E5-1660 3.20 GHz CPU.

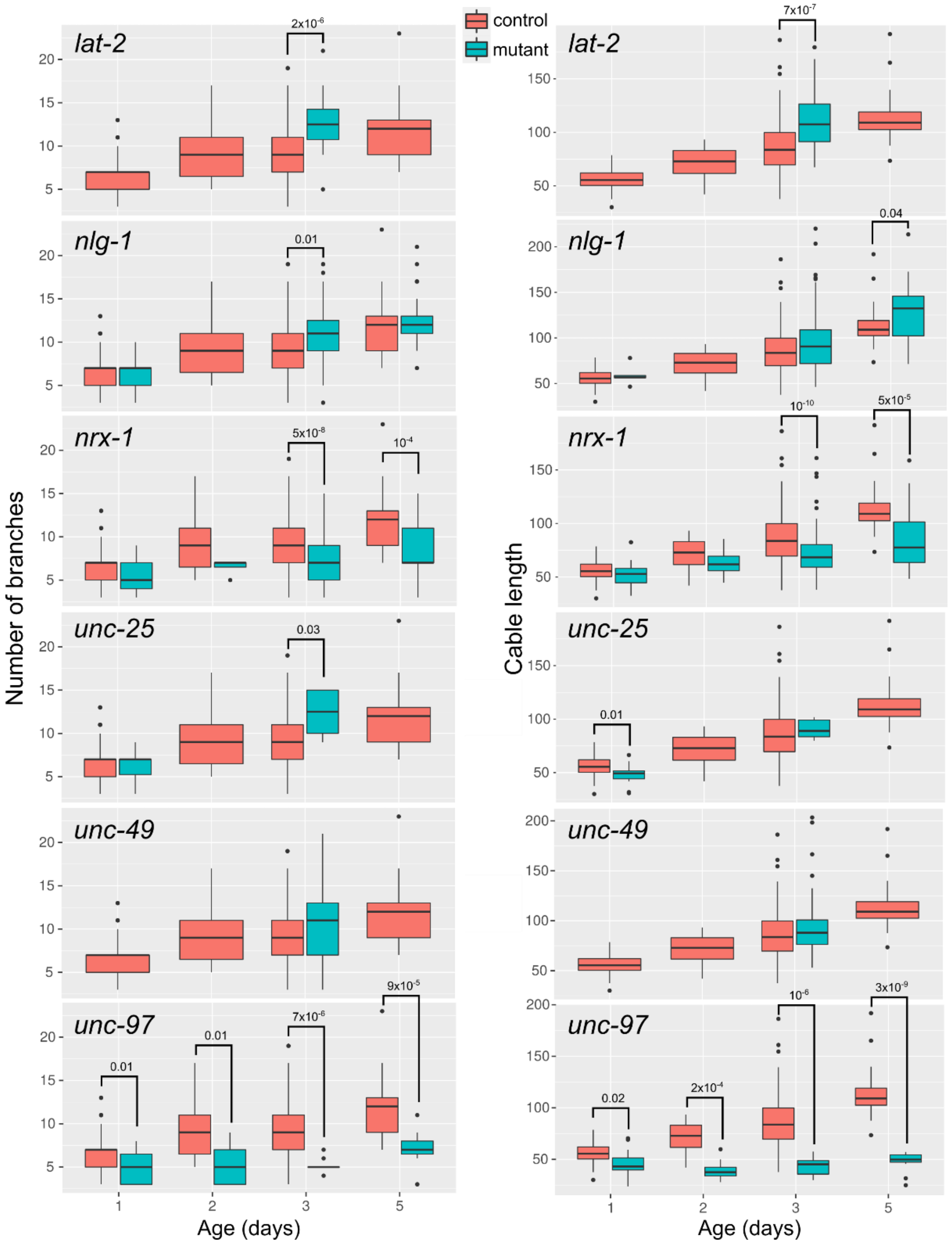


**Supplementary Figure 3. Number of branches and cable length of the DVB neuron in control and mutant worms.** The number of branches and total cable length is shown for each of the 6 significantly associated genes with the morphology of the DVB neuron that were identified with *CAJAL*.

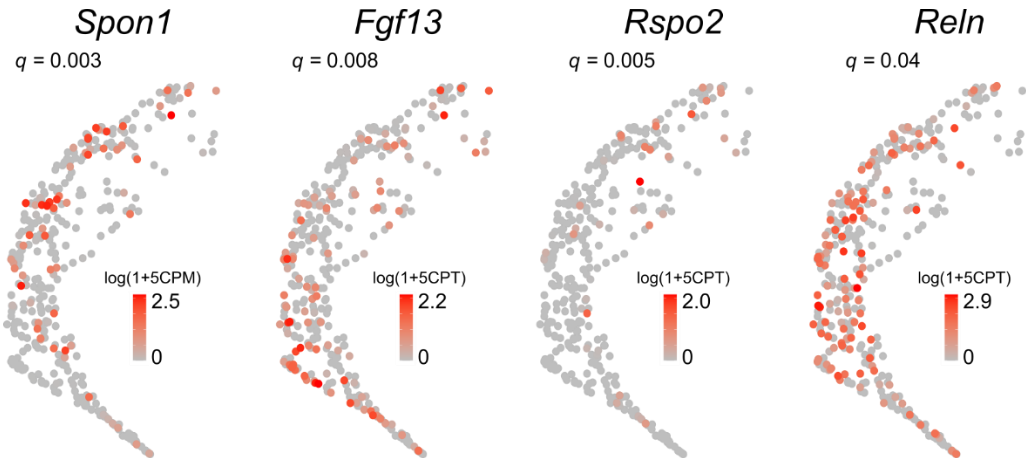


**Supplementary Figure 4. Secreted factors associated with morpho-transcriptomic trajectories of inhibitory neurons.** The UMAP representation of the cell morphology space of the dendrites of 370 inhibitory neurons is colored by the gene expression level of 4 genes coding for secreted factors that are significantly associated with both the RNA velocity field and the structure of the cell morphology space (Laplacian score permutation test, FDR < 0.05). The *q*-value of the Laplacian score is shown for each gene. CPT: counts per thousand.

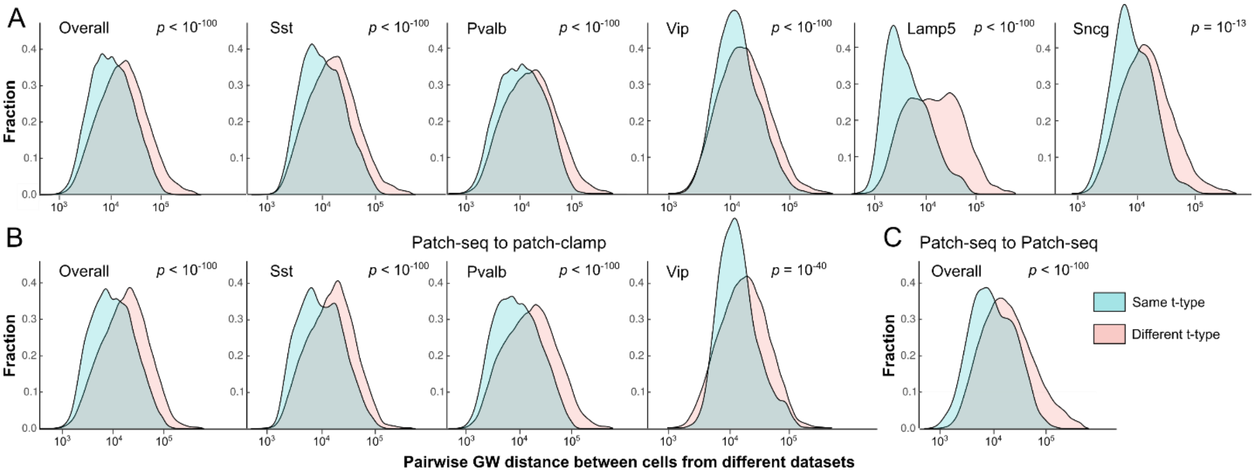


**Supplementary Figure 5. Consistency of major t-types across datasets in the integrated cell morphology space of Patch-seq and patch-clamp datasets. A)** The distributions of pairwise GW distances between cells of the same t-type (blue) and between cells of different t-type (red), where each cell in the pair belongs to a different dataset, are shown for the combined morphology space of the basal and apical dendrites of visual cortex neurons profiled with patch-clamp (Gouwens et al., 2019), and visual cortex and motor cortex neurons profiled with Patch-seq (Gouwens et al., 2020; Scala et al., 2021). In addition to the overall distributions, the distributions restricted to cells from each major t-type are shown. The Wilcoxon rank-sum test *p*-value is indicated in each case. **B)** Same as in (A) but restricting to only distances between Patch-seq and patch-clamp cells. **C)** Same as in (A) but restricting to only distances between cells from the two Patch-seq datasets.

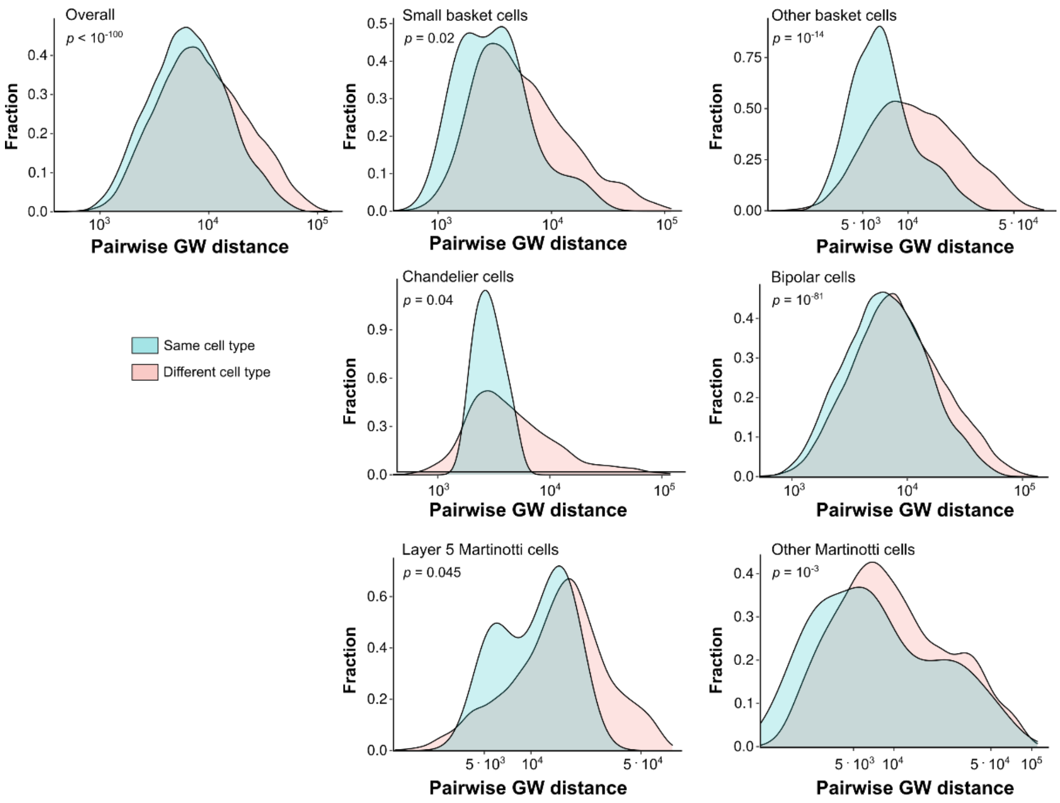


**Supplementary Figure 6. Consistency of cell types across datasets in the integrated cell morphology space of Patch-seq and MICrONS datasets.** The distributions of pairwise GW distances between cells of the same type (blue) and between cells of different type (red), where one cell in the pair belongs to the MICrONS dataset and the other to one of the Patch-seq datasets, are shown for the full reconstructions visual cortex and motor cortex neurons profiled with Patch-seq (Gouwens et al., 2020; Scala et al., 2021) and visual cortical neurons profiled with serial electron microscopy by the MICrONS program (MICrONS Consortium et al., 2021). Cell types in the Patch-seq datasets were annotated based on their t-type. In addition to the overall distributions, the distributions restricted to cells from each type are shown. The Wilcoxon rank-sum test *p*-value is indicated in each case.

### Supplementary Tables

**Supplementary Table 1. *C. elegans* strains used in the morphological analysis of the DVB neuron.** [Table provided as a separate file].

**Supplementary Table 2. Morphological features associated with the structure of the cell morphology space of the DVB neuron.** The Laplacian score, *p*-value, and *q*-value are presented for each of 33 morphological features evaluated in the cell morphology space of the DVB neuron. [Table provided as a separate file].

**Supplementary Table 3. Genes associated with the structure of the cell morphology space of the basal and apical dendrites motor cortex neurons profiled with Patch-seq.** The Laplacian score, *p*-value, and *q*-value are presented for the expression of each gene evaluated in the cell morphology spaces of excitatory and inhibitory neurons. [Table provided as a separate file].

**Supplementary Table 4. Electrophysiological features associated with the structure of the cell morphology space of the basal and apical dendrites motor cortex neurons profiled with Patch-seq.** The Laplacian score, *p*-value, and *q*-value are presented for each of 29 electrophysiological features evaluated in the cell morphology spaces of excitatory and inhibitory neurons. [Table provided as a separate file].

**Supplementary Table 5. Genes associated with morpho-transcriptomic trajectories of inhibitory motor cortex neurons profiled with Patch-seq.** The Laplacian score, *p*-value, and *q*-value evaluated in the cell morphology space are presented for each of the 78 genes associated with the RNA velocity field. [Table provided as a separate file].
